# Inferring time-dependent migration and coalescence patterns from genetic sequence and predictor data in structured populations

**DOI:** 10.1101/342329

**Authors:** Nicola F. Müller, Gytis Dudas, Tanja Stadler

## Abstract

Population dynamics can be inferred from genetic sequence data using phylodynamic methods. These methods typically quantify the dynamics in unstructured populations or assume the parameters describing the dynamics to be constant through time in structured populations. Inference methods allowing for structured populations and parameters to vary through time involve many parameters which have to be inferred. Each of these parameters might be however only weakly informed by data. Here we introduce an approach that uses so-called predictors, such as geographic distance between locations, within a generalized linear model to inform the population dynamic parameters, namely the time-varying migration rates and effective population sizes under the marginal approximation of the structured coalescent. By using simulations, we show that we are able to reliably infer the parameters from phylogenetic trees. We then apply this framework to a previously described Ebola virus dataset. We infer incidence to be the strongest predictor for effective population size and geographic distance the strongest predictor for migration. This allows us to show not only on simulated data, but also on real data, that we are able to identify reasonable predictors. Overall, we provide a novel method that allows to identify predictors for migration rates and effective population sizes and to use these predictors to quantify migration rates and effective population sizes. Its implementation as part of the BEAST2 software package MASCOT allows to jointly infer population dynamics within structured populations, the phylogenetic tree, and evolutionary parameters.

## Introduction

Genetic sequence data can be used to reconstruct the shared evolutionary history or phylogenetic tree of pathogens. These trees are shaped by migration and transmission dynamics, which in turn can be quantified from these trees by using so-called phylodynamic methods. These methods however typically assume that all sequences are from the same well-mixed population.

Methods that account for population structure such as the structured coalescent (Takahata, 1988; Hudson, 1990; Notohara, 1990) allow to infer how lineages coalesce within sub-populations and migrate between them. This is done by inferring effective population sizes and migration rates, with the effective population sizes being related to transmission dynamics (Volz <u>et al.,</u> 2009a). Even when only considering constant parameters through time, the number of parameters to estimate grows quadratically with the number of sub-populations. When additionally considering these parameters to change through time using mathematically tractable skyline approaches, this number has to be multiplied by the number of time points considered. The information in a single phylogenetic tree can however be too limited to inform all these parameters. This problem is not only limited to structured coalescent process but also applies to structured birth-death models (Stadler and Bonhoeffer, 2013; Kühnert <u>et al.,</u> 2016).

Alternatively, one can make use of additional data, such as transportation data, that potentially predict these parameters by using so-called generalized linear models (GLM) (Lemey <u>et al.,</u> 2014). This has for example been done to study the cross-species transmission of bat rabies virus (Faria <u>et al.,</u> 2013) or spatial spread of dog rabies virus through rural Tanzania (Brunker <u>et al.,</u> 2017). It was further used to study Ebola virus dissemination throughout West Africa (Dudas <u>et al.,</u> 2017) or the spread of Dengue virus in the Americas (Nunes <u>et al.,</u> 2014). It has further been extended to inform effective population sizes through time in unstructured populations (Gill <u>et al.,</u> 2016)

The underlying migration model (Lemey <u>et al.,</u> 2009) used in these structured models, although computationally feasible, relies on simplifying assumptions of the tree generating process. Namely, it is assumed that the process which generated the phylogenetic tree is independent from the migration process. This essentially assumes that any two lineages have the same probability of coalescing, no matter where they are. This in turn can lead to biased estimates of migration rates, for example when sampling is biased (De Maio <u>et al.,</u> 2015). Additionally, since only the migration process is modelled, information about the coalescent process in different sub-populations can not be incorporated.

The structured coalescent (Takahata, 1988; Hudson, 1990; Notohara, 1990) on the other hand does not make this independence assumption. This enables us to model the tree generating process by using coalescence within and migration between sub-populations. It however only allows a very limited number of different sub-populations to be considered (Vaughan <u>et al.,</u> 2014), due to computational issues (De Maio <u>et al.,</u> 2015).

In order to allow for structured models with many different sub-populations and parameters that change through time, alternative approaches have been developed (Volz, 2012). By avoiding the sampling of migration histories by formally integrating over every possible migration history, these approaches allow to consider scenarios with more parameters (De Maio <u>et al.,</u> 2015). These approaches have however been subject to strong biases due to simplifying assumptions that were initially not being accounted for (Müller <u>et al.,</u> 2017). The marginal approximation of the structured coalescent on the other hand allows to integrate over every possible migration history, avoiding such biases (Müller <u>et al.,</u> 2017, 2018).

We here introduce an approach based on this approximation that infers the time varying effective population sizes of and the migration rates between different sub-populations from predictor and sequence data, with predictor data icharacterizing one location (e.g. population size, location) or the interaction of two locations (e.g. transportation, distance). We do so by using a generalized linear model where we infer to what degree each predictor predicts effective population sizes or migration rates. By using simulations, we show that we are able to retrieve the extent to which each predictor informs the population dynamics parameters. We then apply our GLM approach to a subset of sequence data from the West African Ebola virus (EBOV) dataset (Dudas <u>et al.,</u> 2017). This subset is comprised of lineages descended from the major introduction of EBOV into Sierra Leone (Dudas <u>et al.,</u> 2017), further downsampled to only sequences collected in 2014. The Sierra Leonean lineage was sustained via intense endemic transmission, making it the dominant EBOV lineage in the entire epidemic (Dudas <u>et al.,</u> 2017). Following its introduction into Sierra Leone this lineage was also the source of EBOV in neighbouring Liberia and Guinea in the late stages of the epidemic (Dudas <u>et al.,</u> 2017). Using the example of Ebola, we demonstrate that our approach is able to retrieve reasonable predictors for the migration rates and effective population sizes.

### New Approaches

The marginal approximation of the structured coalescent (Müller <u>et al.,</u> 2017) allows to consider datasets with many different sub-populations. When one wants to consider time varying rates alongside many different sub-populations, each individual rate in a given time interval is potentially only weakly informed by the phylogentic tree.

To inform rates, Lemey <u>et al.</u> (2014) introduced a generalized linear model approach to estimate migration between different sub-populations. This allows to inform these rates not only from the phylogenetic tree, but also from predictor data. We here apply this generalized linear model approach to inform effective population sizes and migration rates that vary through time.

Similar to (Lemey <u>et al.,</u> 2014), we define the effective population sizes and migration rates as log-linear combinations of coefficients, indicators and time varying predictors. These predictors describe differences of effective population sizes of different sub-populations or migration rate differences between sub-populations. Some previously used predictors for migration rates include air traffic data between different locations (Lemey <u>et al.,</u> 2014; Nunes <u>et al.,</u> 2014) and distances between them (Dudas <u>et al.,</u> 2017). The indicators and coefficients quantify if and to what degree each predictor contributes in predicting effective population size or migration rate differences across different sub-populations and points in time. Whereas the use of indicators is not strictly necessary, they allow us to use priors on the number of active predictors, thereby helping to reduce over-fitting. We implemented this approach as part of the BEAST2 (Bouckaert <u>et al.,</u> 2012) package MASCOT (Müller <u>et al.,</u> 2018). This allows to co-infer indicators and coefficients from genetic sequence and predictor data alongside phylogenetic trees and evolutionary parameters.

## Results

### Inference of predictor contributions from phylogenetic trees

We first tested how well indicators and coefficients can be inferred from phylogenetic trees. We randomly simulated 20 time-varying migration rate and 20 time-varying effective population size predictors. Each value of each predictor at every point in time was drawn from a normal distribution with mean=0 and sigma=1. As in (Lemey <u>et al.,</u> 2014), we standardized each predictor to have mean 0 and standard deviation 1. Next, we randomly chose 3 of the 20 migration rate and 3 of the 20 effective population size predictors to be active predictors. As such, they actually predict migration rate and effective population sizes. All other predictors are considered inactive and are used only to see if inactive predictors can be reliably identified as such. Each of the active predictors was then assigned a random coefficient from a normal distribution with mean=0 and sigma=0.5. By using equations 1 and 2, we then calculated the migration rates between every sub-population and the effective population size in every sub-population at any point in time from the active predictors and coefficients. We then used these parameters to simulate phylogenetic trees using MASTER (Vaughan and Drummond, 2013) under the exact structured coalescent with 1000 serially sampled lineages. Next, we inferred which predictors explained patterns of migration and effective population size in simulated phylogenies and their relative contributions (coefficients).

In figure 1, we show the inferred coefficient values of active predictors as well as the probability that active predictors are identified as such. The coefficients are inferred well for both migration rate and effective population size predictors. While inactive predictors are reliably excluded and predictors with strong effects (large coefficients) are reliably included, predictors with only minor effects (small coefficients) can be falsely excluded. This is however expected due to a small effect size.

**Figure 1:**
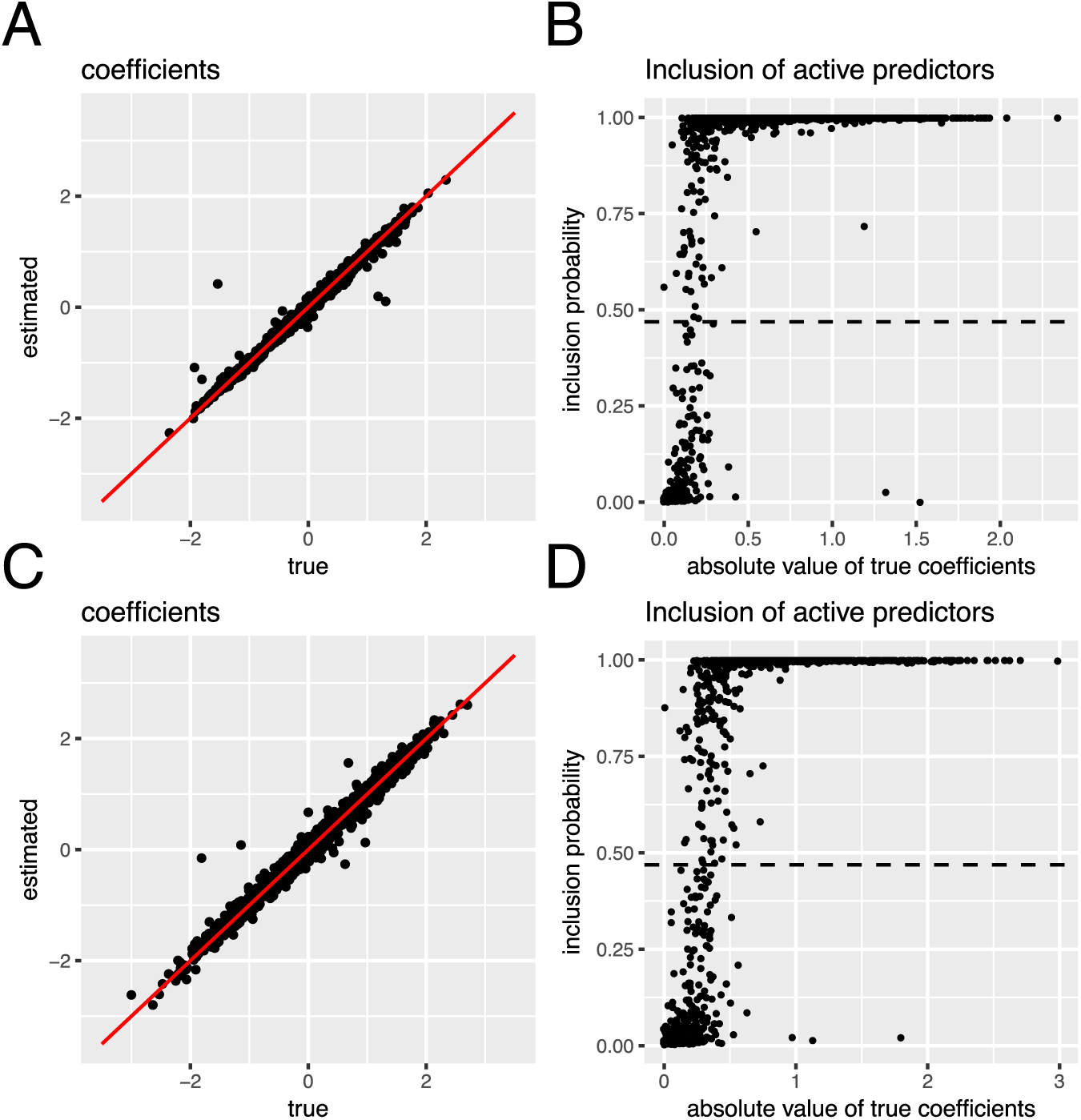
Inference of coefficients and indicators from phylogenetic trees. **A** Inferred active coefficients (y-axis) versus the true coefficients for the effective population size predictors. **B** Probability of the indicator of active predictors to be 1 (y-axis) for the effective population size. The dashed line indicates inclusion probabilities that are equivalent to a Bayes Factors of 5. **C** Inferred active coefficients (y-axis) versus the true coefficients for the migration rate predictors. **D** Probability of the indicator of active predictors to be 1 (y-axis) for the migration rate.

### 2014 Ebola epidemic in Sierra Leone

We used EBOV sequences sampled in 14 different regions of Sierra Leone in 2014 (Dudas <u>et al.,</u> 2017). As migration rate predictors, we used the same time invariant predictors as Dudas <u>et al.</u> (2017). Namely, we used mean travel time to the nearest major settlement of at least 100,000 inhabitants, gridded economic output, population size and density, mean annual temperature and precipitation, and index of precipitation and temperature seasonality. All these predictors can either inform migration rates from or to a particular county and are therefore called origin/destination predictors. Additionally, we added predictors to account for possible random effects of migration between counties. Furthermore, we used the distances between the different counties as a migration rate predictor. For effective population sizes we used origin/destination predictors from Dudas <u>et al.</u> (2017). These predictors were not used previously, since information about the coalescent process in different sub-populations could not be incorporated in previous approaches Lemey <u>et al.</u> (2014).

Additionally, we incorporated the weekly case data of each location as a time variant predictor of the effective population size. Instead of using 0 for weeks with no reported incidences, we used 0.01 in order to not completely exclude lineages to be in a location if there are no reported cases there. This also avoids computational issues arising for effective population sizes being 0 and therefore coalescent rates being 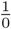. Figure 2 shows the inferred maximum clade credibility tree with the different colors denoting the inferred locations of the nodes calculated as shown in Müller <u>et al.</u> (2018). Further, predictors for the migration rates and the effective population sizes with a Bayes Factor of more than 5 are shown. Incidence is inferred to be the strongest predictor of effective population size. This is to be expected since incidence should be approximately proportional to viral effective population sizes in each location given similarity in transmission rates (Volz <u>et al.,</u> 2009b).

**Figure 2:**
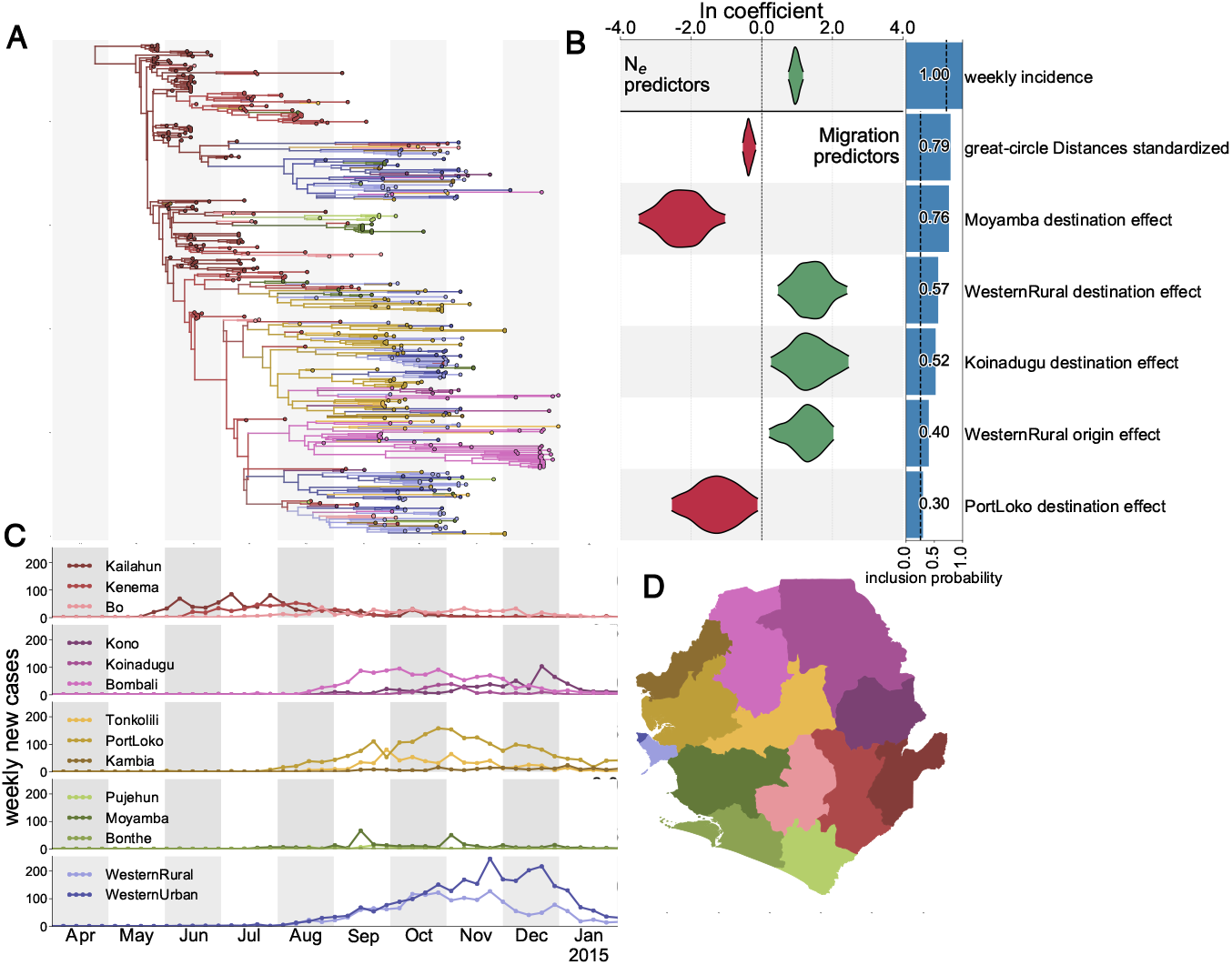
2014 Ebola epidemic in Sierra Leone. **A** Inferred maximum clade credibility tree from the 2014 Sierra Leone EBOV sequences. Colors denote the most likely inferred district for each node, and branches are coloured as their descendant node. District color scheme is shown on the map. **B** Inferred migration rate and effective population size predictors. Violin plots show estimates of coefficients for predictors with a Bayes Factor of more than 5. Bar plots show the mean value of indicators for predictors with BF> 5. **C** Weekly incidence by district. The x-axis denotes time in months and acts as a scale for both incidence data as well as the phylogenetic tree. **D** Color map of districts in Sierra Leone.

We infer great circle distances between different locations to be the strongest predictor for migration rates. This means that migration rates are inferred the strongest between regions whose population centroids are closer. The root of the tree is inferred to be in Kailahun with a 63% probability, with the rest of the probability mass approximately evenly distribution across the other locations. The 63% is possibly overestimated due to not considering un-sampled locations outside Sierra Leone.

## Discussion

We here introduce a method that is able to infer time varying predictors of effective population sizes and migration rates. Previous GLM approaches were restricted to time invariant (Lemey <u>et al.,</u> 2014) and time variant (Bielejec <u>et al.,</u> 2014) rates in models that did not allow to jointly consider the migration and coalescent process. Other approach were using the GLM framework to inform effective population sizes through time in unstructured populations (Gill <u>et al.,</u> 2016). In contrast to that, we here allow to model the migration and coalescent process jointly. While other approach that allow to infer effective population sizes from phylogenetic trees and incidence jointly in structured populations exist (Rasmussen <u>et al.,</u> 2011, 2014), they currently are computationally not feasible for larger datasets.

By using simulations, we show that indicators and coefficients of predictors can be inferred reliably. Predictors that do not explain migration rates or effective population sizes are reliably excluded. This however also applies to predictors with small effect size. These are often inferred to not predict effective population sizes or migration rates at all. In contrast to for example Gill <u>et al.</u> (2016), we currently not allow for error terms in the GLM equation. We therefore essentially assume that all or a subset of the predictors fully explain the migration rates and the effective population sizes through time. Future improvements could fill that gap by allowing for such error terms. This in turn would however require efficient operators to sample these. Further, it would require to develop reasonable priors on these error terms, similar to the ones used for skyline methods (see for example Drummond <u>et al.</u> (2005) or Minin <u>et al.</u> (2008)). Furthermore, the here presented GLM approach could be applied to inform birth, death, migration and sampling rates through time for structured birth-death models (Stadler and Bonhoeffer, 2013; Kühnert <u>et al.,</u> 2016).

By using the example of the 2014 Sierra Leone Ebola virus disease outbreak, we show that out approach is able to infer the effect size of predictors reasonably from real data as well. We infer weekly case numbers to predict effective population sizes best. For migration rate predictors, the distances between population-weighted centres of different locations is inferred to be the strongest predictor. Previously, distances have been identified as an important predictor of geographic spread for Ebola virus in West Africa, by both phylodynamic (Dudas <u>et al.,</u> 2017) and epidemiological approaches (Kramer <u>et al.,</u> 2016), even when only Sierra Leone is considered (Gustafson and Proctor, 2017). Overall, we infer similar migration rate predictors as Dudas <u>et al.</u> (2017) which used the panmictic and time invariant model described in (Lemey <u>et al.,</u> 2014). We mainly expect there to be great differences in the inference of migration rate predictors between the two approaches when sampling is strongly biased (De Maio <u>et al.,</u> 2015).

Sampling of Ebola cases was fairly dense during the outbreak. Whilst Ebola virus sequencing in West Africa has generally kept up well with increasing numbers of cases (Dudas <u>et al.,</u> 2017), numerous locations are however known to have been under-sampled or un-sampled altogether. For example, an EBOV lineage established early in Conakry prefecture of Guinea resurfaced at least three times during the epidemic (Carroll <u>et al.,</u> 2015; Simon-Loriere <u>et al.,</u> 2015; Quick <u>et al.,</u> 2016). This suggests the presence of a substantial, yet cryptic, localised transmission chain not seen outside of Conakry. It remains unclear how to treat entirely un-sampled locations and what their effects might be on internal node state reconstruction or inference of predictor importance. Future research will need to study these effects of so-called ghost states (Slatkin, 2005) on the generalized linear model approach.

Overall, this newly introduced method allows to include predictor data, such as transportation or incidence data, into phylodynamic analyses. This allows us to infer population dynamic parameters as well as the location of ancestral nodes more reliably in a computationally tractable way. Predictor data, such as the movement of people using mobile phone data (Deville <u>et al.,</u> 2014; Wesolowski <u>et al.,</u> 2015) or the social mixing of different age groups (Mossong <u>et al.,</u> 2008), is increasingly being gathered. This in turn means that methods that are able to combine various sources of information in a computationally feasible way will be playing an ever increasing role in epidemiology.

## Methods and Material

### Effective Population sizes and migration rates as generalized linear models

Instead of inferring the effective population size *Ne_a_*(*t*) of state a at time *t* directly, we define it as a linear combination of *c* different predictors 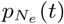, coefficients 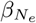 and indicators 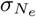:
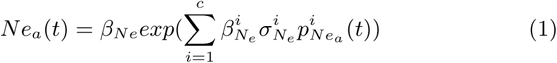

The coefficients 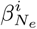 can be between –∞ and ∞ and denote the extent to which each predictor contributes in predicting effective population sizes. The indicators 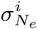 can be 0 or 1 and denote if a predictor contributes at all. This allows to put prior distributions on the number of predictors that are actually used and reduces the issue of over-fitting effective population sizes with too many predictors. We therefore only have to infer the coefficients 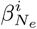 for predictors where the indicator 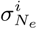 is 1. 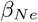 denotes a scaling parameter, scaling every effective population size at every point in time with the same value. The different predictors are in log space and in order to have comparable predictors, they are typically standardized, such that their mean is 0 and their standard deviation is 1. The values of these predictors vary across different states a as well as different time points *t*. This parametrization of the generalized linear model is the same as described in (Lemey <u>et al.,</u> 2014).

We apply the same framework for the forward-in-time varying migration rates 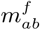 between state *a* and state *b*:
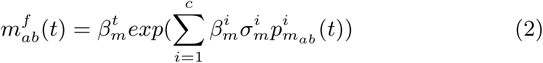

where *β_m_* is the overall rate scaler, describing the overall magnitude of migration. Since the structured coalescent uses backwards in time migration rates, we define the backwards in time rates as:

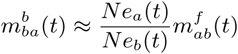

This equality is exact for the case when *α_a_* = *α_b_* such that *Ne_a_*(*t*) = *α_a_I_a_*(*t*) and *Ne_b_*(*t*) = *α_b_I_b_*(*t*) with *I_α_*(*t*) denoting the number of infected in population *a* at time *t*. (Volz, 2012).

### Ebola sequence and incidence data

Sequences belonging to the major Sierra Leonean Ebola virus lineage that dominated the country’s epidemic (Dudas <u>et al.,</u> 2017) were extracted and down-sampled to sequences collected up to 31st of December 2014, leaving 473 taxa. Stretches of putative hypermutation tracts corresponding to hypothesised ADAR edits were identified and masked as described in Dudas <u>et al.</u> (2017).

Incidence data were compiled from the latest WHO report on Ebola virus disease (EVD) cases in Sierra Leone: http://apps.who.int/gho/data/view.ebola-sitrep.ebola-countrySLE-new-conf-prob-districs-20160511-data?lang=en. These data report the number of new EVD cases for each subnational division of Sierra Leone (district) and epi week, split by whether the cases are confirmed or probable. Additionally, due to the scale of the epidemic across the region, there are two databases (an earlier patient database and later situation reports) for EVD incidence that overlap by around a year (2014 Sep - 2015 Sep) with slightly different reported incidences. Available data are likely underestimates of the true burden of EVD in Sierra Leone, and thus we combine confirmed and probable cases, and keep the higher number for each epi week for when the reporting of patient and situation report databases overlap (Dudas <u>et al.,</u> 2017)

### Software

The method above is implemented into our BEAST2 package MASCOT (Marginal Approximation of the Structured COalsescenT). Simulations were performed using a backwards in time stochastic simulation algorithm of the structured coalescent process using MASTER 5.0.2 (Vaughan and Drummond, 2013) and BEAST 2.4.6 (Bouckaert <u>et al.,</u> 2014). Script generation and post-processing were performed in Matlab R2015b. Plotting was done in R 3.2.3 using ggplot2 (Wickham, 2009). Plotting of the EBOV analys was done by using baltic (https://github.com/blab/baltic) and matplotlib (Hunter, 2007). Effective sample sizes for MCMC runs were calculated using coda 0.18-1 (Plummer <u>et al.,</u> 2006).

## Data availability

The source code of the BEAST 2 package MASCOT and the GLM method is available at https://github.com/nicfel/Mascot.git. All scripts for performing the simulations and analyses presented in this paper are available at https://github.com/nicfel/GLM-Material.git. Output files from these analyses, which are not on the github folder, are available upon request from the authors.

## Acknowledgement

We would like to thank Oliver Pybus for his helpful discussion on generalized linear models. We also would like to thank Alexei Drummond and Jing Yang for their helpful comments on the implementation of the generalized linear model. NM and TS are funded in part by the Swiss National Science foundation (SNF; grant number CR32I3_166258). TS is supported in part by the European Research Council under the Seventh Framework Programme of the European Commission (PhyPD: grant agreement number 335529). GD is supported by the Mahan postdoctoral fellowship from the Fred Hutchinson Cancer Research Center and NIH R35 GM119774-01.

## References

Bielejec, F., Lemey, P., Baele, G., Rambaut, A., and Suchard, M. A. (2014). Inferring heterogeneous evolutionary processes through time: from sequence substitution to phylogeography. Systematic biology, 63(4), 493–504.

Bouckaert, R., Lemey, P., Dunn, M., Greenhill, S. J., Alekseyenko, A. V., Drummond, A. J., Gray, R. D., Suchard, M. A., and Atkinson, Q. D. (2012). Mapping the Origins and Expansion of the Indo-European Language Family. Science, 337(6097).

Bouckaert, R., Heled, J., Kühnert, D., Vaughan, T., Wu, C.-H., Xie, D., Suchard, M. A., Rambaut, A., and Drummond, A. J. (2014). BEAST 2: a software platform for Bayesian evolutionary analysis. PLoS computational biology, 10(4), e1003537.

Brunker, K., Lemey, P., Marston, D. A., Fooks, A. R., Lugelo, A., Ngeleja, C., Hampson, K., and Biek, R. (2017). Landscape attributes governing local transmission of an endemic zoonosis: Rabies virus in domestic dogs. Molecular Ecology.

Carroll, M. W., Matthews, D. A., Hiscox, J. A., Elmore, M. J., Pollakis, G., Rambaut, A., Hewson, R., Garcia-Dorival, I., Bore, J. A., Koundouno, R., Abdellati, S., Afrough, B., Aiyepada, J., Akhilomen, P., Asogun, D., Atkinson, B., Badusche, M., Bah, A., Bate, S., Baumann, J., Becker, D., Becker-Ziaja, B., Bocquin, A., Borremans, B., Bosworth, A., Boettcher, J. P., Cannas, A., Carletti, F., Castilletti, C., Clark, S., Colavita, F., Diederich, S., Donatus, A., Duraffour, S., Ehichioya, D., Ellerbrok, H., Fernandez-Garcia, M. D., Fizet, A., Fleischmann, E., Gryseels, S., Hermelink, A., Hinzmann, J., Hopf-Guevara, U., Ighodalo, Y., Jameson, L., Kelterbaum, A., Kis, Z., Kloth, S., Kohl, C., Korva, M., Kraus, A., Kuisma, E., Kurth, A., Liedigk, B., Logue, C. H., Lüdtke, A., Maes, P., McCowen, J., Mély, S., Mertens, M., Meschi, S., Meyer, B., Michel, J., Molkenthin, P., Muñoz-Fontela, C., Muth, D., Newman, E. N. C., Ngabo, D., Oestereich, L., Okosun, J., Olokor, T., Omiunu, R., Omomoh, E., Pallasch, E., Pályi, B., Portmann, J., Pottage, T., Pratt, C., Priesnitz, S., Quartu, S., Rappe, J., Repits, J., Richter, M., Rudolf, M., Sachse, A., Schmidt, K. M., Schudt, G., Strecker, T., Thom, R., Thomas, S., Tobin, E., Tolley, H., Trautner, J., Vermoesen, T., Vitoriano, I., Wagner, M., Wolff, S., Yue, C., Capobianchi, M. R., Kretschmer, B., Hall, Y., Kenny, J. G., Rickett, N. Y., Dudas, G., Coltart, C. E. M., Kerber, R., Steer, D., Wright, C., Senyah, F., Keita, S., Drury, P., Diallo, B., de Clerck, H., Van Herp, M., Sprecher, A., Traore, A., Diakite, M., Konde, M. K., Koivogui, L., Magassouba, N., Avšič-Županc, T., Nitsche, A., Strasser, M., Ippolito, G., Becker, S., Stoecker, K., Gabriel, M., Raoul, H., Di Caro, A., Wölfel, R., Formenty, P., and Günther, S. (2015). Temporal and spatial analysis of the 2014–2015 Ebola virus outbreak in West Africa. Nature, 524(7563), 97–101.

De Maio, N., Wu, C.-H., O’Reilly, K. M., and Wilson, D. (2015). New Routes to Phylogeography: A Bayesian Structured Coalescent Approximation. PLoS genetics, 11(8), e1005421.

Deville, P., Linard, C., Martin, S., Gilbert, M., Stevens, F. R., Gaughan, A. E., Blondel, V. D., and Tatem, A. J. (2014). Dynamic population mapping using mobile phone data. Proceedings of the National Academy of Sciences, 111(45), 15888–15893.

Drummond, A. J., Rambaut, A., Shapiro, B., and Pybus, O. G. (2005). Bayesian coalescent inference of past population dynamics from molecular sequences. Molecular biology and evolution, 22(5), 1185–1192.

Dudas, G., Carvalho, L. M., Bedford, T., Tatem, A. J., Baele, G., Faria, N. R., Park, D. J., Ladner, J. T., Arias, A., Asogun, D., et al. (2017). Virus genomes reveal factors that spread and sustained the ebola epidemic. Nature, 544(7650), 309–315.

Faria, N. R., Suchard, M. A., Rambaut, A., Streicker, D. G., and Lemey, P. (2013). Simultaneously reconstructing viral cross-species transmission history and identifying the underlying constraints. Philosophical Transactions of the Royal Society of London B: Biological Sciences, 368(1614), 20120196.

Gill, M. S., Lemey, P., Bennett, S. N., Biek, R., and Suchard, M. A. (2016). Understanding past population dynamics: Bayesian coalescent-based modeling with covariates. Systematic biology, 65(6), 1041–1056.

Gustafson, K. B. and Proctor, J. L. (2017). Identifying spatio-temporal dynamics of Ebola in Sierra Leone using virus genomes. Journal of The Royal Society Interface, 14(136), 20170583.

Hudson, R. R. (1990). Gene genealogies and the coalescent process. Oxford surveys in evolutionary biology, 7(1), 44.

Hunter, J. D. (2007). Matplotlib: A 2d graphics environment. Computing In Science & Engineering, 9(3), 90–95.

Kramer, A. M., Pulliam, J. T., Alexander, L. W., Park, A. W., Rohani, P., and Drake, J. M. (2016). Spatial spread of the West Africa Ebola epidemic. Royal Society Open Science, 3(8), 160294.

Kühnert, D., Stadler, T., Vaughan, T. G., and Drummond, A. J. (2016). Phylodynamics with migration: a computational framework to quantify population structure from genomic data. Molecular biology and evolution, 33(8), 2102–2116.

Lemey, P., Rambaut, A., Drummond, A. J., and Suchard, M. a. (2009). Bayesian phylogeography finds its roots. PLoS Computational Biology, 5(9), e1000520.

Lemey, P., Rambaut, A., Bedford, T., Faria, N., Bielejec, F., Baele, G., Russell, C. A., Smith, D. J., Pybus, O. G., Brockmann, D., and Suchard, M. A. (2014). Unifying viral genetics and human transportation data to predict the global transmission dynamics of human influenza H3N2. PLoS pathogens, 10(2), e1003932.

Minin, V. N., Bloomquist, E. W., and Suchard, M. A. (2008). Smooth skyride through a rough skyline: Bayesian coalescent-based inference of population dynamics. Molecular biology and evolution, 25(7), 1459–1471.

Mossong, J., Hens, N., Jit, M., Beutels, P., Auranen, K., Mikolajczyk, R., Massari, M., Salmaso, S., Tomba, G. S., Wallinga, J., et al. (2008). Social contacts and mixing patterns relevant to the spread of infectious diseases. PLoS medicine, 5(3), e74.

Müller, N. F., Rasmussen, D. A., and Stadler, T. (2017). The structured coalescent and its approximations. Molecular Biology and Evolution, page msx186.

Müller, N. F., Rasmussen, D., and Stadler, T. (2018). Mascot: Parameter and state inference under the marginal structured coalescent approximation. Bioinformatics, page bty406.

Notohara, M. (1990). The coalescent and the genealogical process in geographically structured population. Journal of Mathematical Biology, 29(1), 59–75.

Nunes, M. R. T., Palacios, G., Faria, N. R., Sousa, Jr, E. C., Pantoja, J. A., Rodrigues, S. G., Carvalho, V. L., Medeiros, D. B. A., Savji, N., Baele, G., Suchard, M. A., Lemey, P., Vasconcelos, P. F. C., and Lipkin, W. I. (2014). Air travel is associated with intracontinental spread of dengue virus serotypes 1–3 in brazil. PLOS Neglected Tropical Diseases, 8(4), 1–13.

Plummer, M., Best, N., Cowles, K., and Vines, K. (2006). Coda: convergence diagnosis and output analysis for mcmc. R News, 6(1), 7–11.

Quick, J., Loman, N. J., Duraffour, S., Simpson, J. T., Severi, E., Cowley, L., Bore, J. A., Koundouno, R., Dudas, G., Mikhail, A., Ouédraogo, N., Afrough, B., Bah, A., Baum, J. H. J., Becker-Ziaja, B., Boettcher, J. P., Cabeza-Cabrerizo, M., Camino-Sánchez, A., Carter, L. L., Doerrbecker, J., Enkirch, T., Dorival, I. G., Hetzelt, N., Hinzmann, J., Holm, T., Kafetzopoulou, L. E., Koropogui, M., Kosgey, A., Kuisma, E., Logue, C. H., Mazzarelli, A., Meisel, S., Mertens, M., Michel, J., Ngabo, D., Nitzsche, K., Pallasch, E., Patrono, L. V., Portmann, J., Repits, J. G., Rickett, N. Y., Sachse, A., Singethan, K., Vitoriano, I., Yemanaberhan, R. L., Zekeng, E. G., Racine, T., Bello, A., Sall, A. A., Faye, O., Faye, O., Magassouba, N., Williams, C. V., Amburgey, V., Winona, L., Davis, E., Gerlach, J., Washington, F., Monteil, V., Jourdain, M., Bererd, M., Camara, A., Somlare, H., Camara, A., Gerard, M., Bado, G., Baillet, B., Delaune, D., Nebie, K. Y., Diarra, A., Savane, Y., Pallawo, R. B., Gutierrez, G. J., Milhano, N., Roger, I., Williams, C. J., Yattara, F., Lewandowski, K., Taylor, J., Rachwal, P., J. Turner, D., Pollakis, G., Hiscox, J. A., Matthews, D. A., Shea, M. K. O., Johnston, A. M., Wilson, D., Hutley, E., Smit, E., Di Caro, A., Wölfel, R., Stoecker, K., Fleischmann, E., Gabriel, M., Weller, S. A., Koivogui, L., Diallo, B., Keïta, S., Rambaut, A., Formenty, P., Günther, S., and Carroll, M. W. (2016). Real-time, portable genome sequencing for Ebola surveillance. Nature, 530(7589), 228–232.

Rasmussen, D. A., Ratmann, O., and Koelle, K. (2011). Inference for nonlinear epidemiological models using genealogies and time series. PLoS computational biology, 7(8), e1002136.

Rasmussen, D. A., Volz, E. M., and Koelle, K. (2014). Phylodynamic inference for structured epidemiological models. PLoS computational biology, 10(4), e1003570.

Simon-Loriere, E., Faye, O., Faye, O., Koivogui, L., Magassouba, N., Keita, S., Thiberge, J.-M., Diancourt, L., Bouchier, C., Vandenbogaert, M., Caro, V., Fall, G., Buchmann, J. P., Matranga, C. B., Sabeti, P. C., Manuguerra, J.-C., Holmes, E. C., and Sall, A. A. (2015). Distinct lineages of Ebola virus in Guinea during the 2014 West African epidemic. Nature, 524(7563), 102–104.

Slatkin, M. (2005). Seeing ghosts: the effect of unsampled populations on migration rates estimated for sampled populations. Molecular Ecology, 14(1), 67–73.

Stadler, T. and Bonhoeffer, S. (2013). Uncovering epidemiological dynamics in heterogeneous host populations using phylogenetic methods. Philosophical Transactions of the Royal Society B: Biological Sciences, 368(1614), 20120198–20120198.

Takahata, N. (1988). The coalescent in two partially isolated diffusion populations. Genetical research, 52(3), 213–222.

Vaughan, T. G. and Drummond, A. J. (2013). A stochastic simulator of birth-death master equations with application to phylodynamics. Molecular biology and evolution, 30(6), 1480–93.

Vaughan, T. G., Kühnert, D., Popinga, A., Welch, D., and Drummond, A. J. (2014). Efficient Bayesian inference under the structured coalescent. Bioinformatics (Oxford, England), 30(16), 2272–9.

Volz, E. M. (2012). Complex population dynamics and the coalescent under neutrality. Genetics, 190(1), 187–201.

Volz, E. M., Kosakovsky Pond, S. L., Ward, M. J., Leigh Brown, A. J., and Frost, S. D. W. (2009a). Phylodynamics of infectious disease epidemics. Genetics, 183(4), 1421–30.

Volz, E. M., Pond, S. L. K., Ward, M. J., Brown, A. J. L., and Frost, S. D. (2009b). Phylodynamics of infectious disease epidemics. Genetics, 183(4), 1421–1430.

Wesolowski, A., Metcalf, C., Eagle, N., Kombich, J., Grenfell, B. T., Bjørnstad, O. N., Lessler, J., Tatem, A. J., and Buckee, C. O. (2015). Quantifying seasonal population fluxes driving rubella transmission dynamics using mobile phone data. Proceedings of the National Academy of Sciences, 112(35), 11114–11119.

Wickham, H. (2009). ggplot2: Elegant Graphics for Data Analysis. Springer-Verlag New York.

